# A mathematical model for pancreatic cancer during intraepithelial neoplasia

**DOI:** 10.1101/2024.03.16.585362

**Authors:** Joshua Briones-Andrade, Guillermo Ramírez-Santiago, J. Roberto Romero-Arias

## Abstract

Cancer is the result of complex interactions of intrinsic and extrinsic cell processes, which promote sustained proliferation, resistance to apoptosis, reprogramming and reorganization. To understand the evolution of any type of cancer it is necessary to understand the role of the microenvironmental conditions and the impact of some molecular complexes and mechanisms on certain signalling pathways. As in most cancer quantitative models, the understanding of the early onset of cancer requires a multiscale analysis of the cellular microenvironment. In this paper we analyse a multiscale model of pancreatic adenocarcinoma by modelling the cellular microenvironment through elastic cell interactions and their intercellular communication mechanisms, such as growth factors and cytokines. We focus on the low-grade dysplasia (PanIN 1) and moderate dysplasia (PanIN 2) stages of the pancreatic adenocarcinoma. To this end we propose a gene regulatory network associated with the processes of proliferation and apoptosis of pancreatic cells and its kinetics in terms delayed differential equations to mimic cell development. Likewise, we couple the cell cycle with the spatial distribution of cells and the transport of growth factors to show that the adenocarcinoma evolution is triggered by inflammatory processes. We show that the oncogene RAS may be an important target to develop anti-inflammatory strategies that limit the emergence of more aggressive adenocarcinomas.

## Introduction

Pancreatic ductal adenocarcinoma is among the seventh leading causes of cancer-related deaths wroldwide due to its extreme difficulty to be detected and treated [1, 2]. There are different types of pancreatic cancer, but the most common is adenocarcinoma in about 95% of cases. It originates in exocrine cells in the pancreatic duct lining. Islet cell carcinoma begins in the endocrine cells, and are malignant mostly. Because the pancreas is located deep in the body, a pancreatic tumor is impossible to detect during a standard physician’s examination. What complicates even more the situation is the fact that there are no typical symptoms of a pancreatic tumor until metastasis occurs.

To better understand the key molecular mechanisms of the development of pancreatic cancer, researchers have recently focused on analyzing the role of microenvironment in the development of pancreatic cancer. The microenvironment consists of diverse components that include metabolites, fibroblasts, endothelial cells, immune cells, as well as endocrine cells, that interact with each other together with cancer cells in a complex fashion. This interplay of interactions has important implications for pancreatic cancer cell growth, migration, invasion, angiogenesis, and immunological recognition of cancer cells.

Unlike normal cells, tumor cells often increase their glucose consumption and lactate production even in the presence of physiological oxygen concentrations and functional mitochondria. Solute carriers of the glucose transporter (GLUT) family mediate the first step for cellular glucose consumption. The upregulation of GLUTs has been reported in numerous cancer types as a result of perturbation of gene expression or protein relocalization or stabilization [3]. There are fourteen GLUT proteins that are classified into three classes according to their sequence similarity, namely: Class 1 (GLUTs 1–4, and 14); Class 2 (GLUTs 5, 7, 9, and 11); and Class 3 (GLUTs 6, 8, 10, 12, and 13/HMIT) [3]. Recent investigations have identified GLUT1 and GLUT3 as the main proteins that accelerate cell metabolism [4]. High expression of GLUT1 and/or GLUT3 has been associated with poor patient survival in colorectal carcinoma, breast carcinoma, lung adenocarcinoma, squamous cell carcinoma, ovarian carcinoma, and glioblastoma [5–10]. As discussed in [3], deregulated GLUT1 and GLUT3 expression happens either via a direct interaction with gene regulatory elements, or by controlling the cellular trafficking of the transporters to the membranes.

The microenvironment can be modeled with a multiscale model which is based on biological regulation that consists of complex dynamical interacting biomolecular networks. Transcription factors control the production of other transcription factors and kinases as well as the activation of other kinases and cells behavior. Multiscale modelling of pancreatic carcinoma considers the role of pancreatic cancer cells (PCCs) and pancreatic stellate cells (PSCs), cytokines and growth factors, which are responsible for intercellular communication between the PCCs and PSCs. Major contributors to this microenvironment include immune cells, endothelial cells, nerve cells, lymphocytes, dendritic cells, the extracellular matrix, and stellate cells. It has been found that molecules and cells surrounding the PCCs significantly impact the cancer’s response to therapy. Recent studies have also shown that cancer cells secrete numerous types of cytokines, which play an important role in intracellular communication between PCCs and PSCs [11].

Populations of cells, multiple time scales and diverse size scales, as well as complex nonlinear dynamics in the analysis of gene regulatory networks (GRN) usually yield fixed points, attractors and different phenotypes. Even though a multiscale model would provide more realistic results, one lacks available control methods that can be applied to the analysis of these models. In spite of this, one can built up a Boolean network (BN) approximation of the model to get a kind of calibration or a guide to the continuous multiscale model [12]. With the BN approximation one can carry out a discrete dynamical analysis in search of phenotype targets, fixed points and attractors as well as GRN responses to different inputs. In addition, network motifs are important substructures of biological GRN’s that describe negative feedback, feed-forward regulation and cascades [13–15]. The functioning of these network motifs often depends on emergent properties of many different parameters, especially, delays [16–19]. Delays are crucial to behavior in genetic oscillators as well as in development and disease [14, 16, 20, 21]. They can lead to significant biological changes, for instance, sustained oscillations [17, 19–21].

Pancreatic cancer is a complex and genetically heterogeneous disease. There are many variants of pancreatic cancer and are highly dependent on tissue type –exocrine or endocrine– and cell of origin within the pancreas. More than 95% of pancreatic cancer cases are derived from pancreatic acinar and ductal cell populations and reside within the exocrine gland tissue [22]. Pancreatic ductal adenocarcinoma (PDAC) is believed to form in the epithelial cell layer of the gland tissue. It originates in a stepwise progression from healthy epithelium to precursor lesion to invasive carcinoma and is believed to arise from a combination of acute trauma, chronic inflammation, and spontaneous genetic mutations of oncogenic KRAS and tumor suppressor gene (TSG) [23]. The incurable pancreatic ductal adenocarcinoma (PDAC) is driven by mutations in constitutively active KRAS [24, 25]. Activating mutations of KRAS found in human impair intrinsic GTPase activity of the KRAS protein and can block the interaction between KRAS and GAPs. This leads to constitutive activation of KRAS and persistent stimulation of downstream signalling pathways that drive many of the cancer hallmarks such as: sustained proliferation, metabolic reprogramming, anti-apoptosis, remodelling of the tumour microenvironment, evasion of the immune response, cell migration and metastasis [25, 26]. Oncogenic KRAS reprograms PDAC cells to a highly anti-apoptotic state [27]. Resistance to apoptosis makes PDAC highly resistant to the mitochondrial pattern of apoptosis-regulated cell death. Here it is important to note that KRAS, together with HRAS and NRAS expressed genes belong to the RAS family of proteins that are proto-oncogenes that are frequently mutated in human cancers. These expressed genes are GTPases that function as molecular switches regulating pathways responsible for proliferation and cell survival [28].

There have been found four distinct precursor lesions that vary in their degree of dysplasia and propensity to develop into infiltrating carcinoma [29]: intraductal papillary mucinous neoplasms (IPMN), mucinous cystic neoplasms (MCN), intraductal tubulopapillary neoplasms (ITPN), and pancreatic intraepithelial neoplasms (PanIN) [29]. IPMNs, MCNs, and ITPNs are visible neoplasia that originate in the ducts epithelial cell layer and invade the lumen and surrounding connective tissue. PanINs are microscopic non-invasive lesions (⪅ 5 mm) that are located well inside the small intralobular pancreatic ducts. They constitute the most common precursor lesion and cannot be detected or surgically resected as in the other precursor lesions [30].

PanINs are sub-classified into PanIN 1, PanIN 2 and PanIN 3, lessions according to the degree of architectural changes and cytological abnormalities [31]. PanINs can evolve starting at grade 1 (PanIN 1) and progress through PanIN 2 and PanIN 3 to malignancy [32]. The risk of these lesions forming invasive carcinoma depends on their genetic stability, location and size inside the pancreas. However, it is strongly believed that most PDAC cases originate from PanIN lesions in cells having developed oncogenic KRAS mutations [31, 33]. Taking this into account, here we present and analyse a multiscale quantitative model that mimics the evolution of low-grade dysplasia (PanIN 1) and moderate dysplasia (PanIN 2) stages of a pancreatic adenocarcinoma.

## Materials and methods

### Model

The pancreas is a complex system made up of several separate functional units that regulate two main physiological processes: digestion and glucose metabolism [35]. In the present study, we will focus on the most common pancreatic cancer that involves the exocrine pancreas which is made up of acinar and ductal cells. It originates from the exocrine cells of the lining of the pancreatic duct [36]. The acinar cells that make up the majority of the tissue are arranged in grape-like clusters found at the smaller ends of the branching duct system. The ducts, which supply an enzyme mixture, in turn, form a network of increasing size that culminates in the main pancreatic ducts. The symmetry of the three-dimensional structure of the pancreas ductal and acinar cells allows us to model it as a two-dimensional projection that represents the end of the duct and the beginning of the acinar group cells in the form of U. This approximation allows the numerical analysis of the model in 2D space and validates it by comparing the obtained patterns from the simulations with histological or optical observations obtained in clinical trials. Pancreatic Intraepithelial Neoplasias (PanINs) are microscopic neoplastic lesions in smaller pancreatic ducts and are often associated with lobulocentric atrophy of the pancreas. PanIN 1 is flat or papillary with columnar epithelium and basally oriented, round nuclei. PanIN 2 is papillary with nuclear hyperchromasia, crowding, and pseudo stratification. PanIN 3 is papillary, micropapillary, or cribriform with nuclear pleomorphism, frequent loss of nuclear orientation, and mitosis [37]. During the stages of neoplasia evolution, cell proliferation rates increase and morphological alterations are increasingly divergent in relation to normal channels, suggesting the generation of abnormal cells in their growth. The identification of common mutational profiles in simultaneous lesions has provided evidence of the relationship between PanINs and the pathogenesis of pancreatic adenocarcinoma [35]. Specifically, common mutation patterns in PanIN and adenocarcinoma associated with the RAS oncogene family have been correlated. Thus, activating mutations on the RAS family genes are known to have a remarkable variety of cellular effects, including the induction of proliferation, survival, and invasion through stimulation of various signaling pathways, including the regenerative capacity of cells and the multipotent differentiation capacity. As a consequence, acinar cells can transdifferentiate into ductal cells in the absence of cell division and newly emerging ductal cells can subsequently redifferentiate into stem cells.Some in vitro studies [34, 35, 38] suggest that identifying the genetic alterations of the KRAS gene is not enough to understand the development of neoplasia. It is also necessary to consider the processes of inflammation or stress conditions and tissue damage as well as generalized activation of KRAS [39, 40]. In this sense, it has been shown that the induction of an inflammatory process generates different predispositions of exocrine cells to tumorigenesis. In addition, it has been shown that acinar and ductal cell lesions may affect the probability of tumor formation. Based on the shape and spatial arrangement of the pancreas, one can conclude that cell reproduction in the early stages of pancreas development mainly involves three intertwined dynamics, namely, cell proliferation, inflammatory process, and glucose concentration gradients, whose occurrence provides the spatial information necessary to regulate each cell proliferation rate and ultimately determine its phenotype. Considering that the elastic properties of the tissues are being used to assist in diagnosis of solid pancreatic lesions (SPL) [41–45] our first hypothesis is that the conditions imposed by the extracellular matrix regulate the processes of growth and proliferation. It has been found that in both mice and humans, tissue stiffness increases in proportion to tumor size, indicating that tuning of mechanical stiffness is an ongoing process during tumor progression [41]. For simplicity, here we propose that there exits a potential energy associated to the elastic mechanical field in the ductal and acinar tissues that can be modelled with a second order polynomial in the tissue deformation [46]. With this characterization, tissue local and global elastic properties can be described without difficulty by specific functions of the elastic stress, mechanical pressure as well as other local mechanical forces. Our second hypothesis considers that cells need to consume glucose to proliferate and grow. Because transport of glucose depends on the tissue stiffness, one can assume that spatio-temporal patterns of glucose concentration in the acinar and ductal tissue are shaped by the local elastic field, [12, 47] and elastic tissue relaxation [48–51]. Taking this into account our third hypothesis assumes that at places where cells divide and tissue expands changes are produced in the elastic field which lead to glucose concentration gradients that determine local cell proliferation rate. To model the elastic properties of the pancreas ductal and acinar cells we define a potential function that depends on time and space. The negative gradient of this function yields the mechanical force. Taking advantage of the spatial symmetry of the acinar cells, the tissue is represented by a two-dimensional tessellation that leads to a set of polygonal cells covering the space without any gaps or overlaps. To this end we use Voronoi diagrams obtained from a collection of scattered random points on a Euclidean plane which represent the positions assigned to each biological cell.

### Voronoi diagrams

A Voronoi diagram is a set of points assigned to a limited geometric region of space in the form of a convex polygon. Voronoi cells are currently used in many fields of science, however, it was Honda [52] who first proposed their use for modeling biological cells. To model the pancreas geometric shape and cell growth we define a geometric space domain as follows: 1) We build two regular shapes with points inside two rectangles that are connected by a double semicircular arc. This arrangement mimics a structure like a horseshoe that will help us to simulate the pancreas inflammatory process in the ductal and acinar cells. The extracellular matrix that surrounds the pancreas tissue is represented by a set of equidistantly separated fixed exterior points. 2) Because the points on the edge cannot define a convex polygon the associated spatial cells will not be closed so that they are not considered in the tissue development under consideration. 3) We randomly place N points inside the domain such that all Voronoi cells are closed and connected. To guarantee this, we use the Delaunay triangulation algorithm. Note that spatial cells areas (A_*i*_) vary in size and shape, and that the generated points (r_*i*_) shown in the figure often times do not correspond to the center of mass of the corresponding biological cells 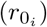. In a homogenization process, the average area 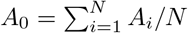 would be the space that each one of the biological cells would occupy. In addition, we observe that the spatial cell distribution forms a regular hexagonal network, which is the network that produces the closest packed structure. In such a case, the distance in the regular array is 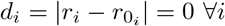. To emulate the inflammation processes we complement this two dimensional geometric model by introducing a elastic potential function 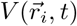 with minima at the equilibrium points configuration 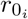.

### Mechanical fields

In most tissues cells mechanically attach to their neighbors through their common interfaces and exert forces upon each other as well as on their surrounding environment. These complex interactions can lead to significant morphogenetic deformations that can lead to alterions in the developing tissue. These deformations can generate crucial folding, stretching or constrictions to establish and define the shape of the organism. Understanding how cells collectively perform these tasks is an important question that lies at the interface of physics, developmental biology, and particularly cancer initiation. Tissue deformation occurs when some mechanical properties of cells change. On the other hand, cell division and apoptosis also impose physical constraints on the surrounding environment. Because of this, it is important to understand the mechanical forces that act on a tissue and how equilibrium is reached. Bearing this in mind we consider the relevant internal and external forces acting on the tissue that balance the friction and viscous forces. Previous models have used springs and vertex models to understand how the complex interplay between cell shape, forces and mechanical constraints are generated within epithelial cells [53] and make epithelial tissues confluent with active matter dynamics as well as complex phenomena arising from other biophysical and mecano-transduction signaling [54–57].

In the present model, we use the average cell area, *A*_0_, as a measurable quantity to identify healthy cells. Any deviation from *A*_0_ will point to an unhealthy cell. In addition, we consider that when cells in the tissue are isotropically shaped a non-zero distance *d*_*i*_ value will ensue as an indication that cells in the tissue are stressed.

To incorporate in the model the elastic properties of cells and tissue a deformation force is introduced by means of a harmonic potential [54] that depends on the cell area *A*_*i*_ and the cell center of mass position, 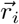.

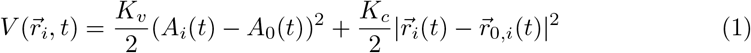

where the coefficients *K*_*v*_ and *K*_*c*_ represent the elastic constants. That is, the elastic potential energy is approximated to second order in the deformation by a Taylor series around the equilibrium biological cell state, 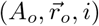.

At this point it is important to recall that biological as well as biophysical systems do not conserve energy. On the contrary, they undergo irreversible processes that consume and dissipate energy [58]. Taking this into account we have also included in the model a dissipation term in the form of a friction force, which for simplicity, we assume it is proportional to biological cell velocity, 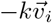. It emulates the biological cell inability to undergo drastic mechanical deformations in shape and/or changes in size. Bearing in mind all the aspects of the above description, one can write down the following equations of motion for the *i − th* biological cell [54]:

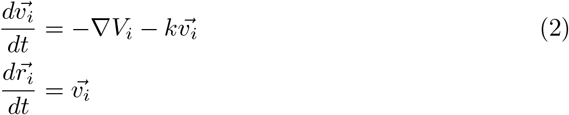

Where 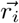 and 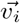 are the position and velocity of the i− th biological cell in the tissue, respectively.

We now turn to the emulation of cell proliferation (cell mitosis) and apoptosis processes. One should recall that the construction of a Voronoi diagram associates a polygonal form to each point of the population [59, 60]. Thus, cell mitosis requires that we choose two points inside a Voronoi spatial cell. In each iteration the Voronoi diagrams are recalculated locally and the physical space is increased to make room for the two daughter cells until they reach the average size A_*i*_. On the other hand, to emulate the apoptosis process a biological cell disappears so that the point associated to it is erased and the Voronoi diagrams are rebuilt locally.

### Glucose transport

Several studies [61–63] suggest that cancer cells development is strongly dependent on glucose consumption since it drives the process of aerobic glycolysis. This in turn generates a greater production of cellular metabolites, which are necessary for the generation of new biomass and nutrients for the diverse signalling processes. Thus, increased glucose metabolism during cancer development is an indication that cancer cells prefer the use of glycolysis as a mean to generate ATP even in the presence of oxygen. By contrast, normal cells metabolize glucose for energy production and resort to the glycolytic pathway under anaerobic conditions [62].

Glucose transport across the plasma membrane is a rate-limiting step in glucose metabolism and is mediated by a family of glucose transporters (GLUTs) [47, 64]. GLUTs allow bidirectional transport of glucose across the cell membrane down its concentration gradient. This metabolic regulation facilitates its incorporation into the biomass of essential nutrients for cell proliferation. There are studies which show that the deregulation of this metabolic pathway in the presence of growth factors, insulin, and stress, leads to alterations in cell differentiation and transformation conditions [62, 65]. It has also been found in different types of cancer that GLUTs are deregulated and can be identified with some cancer expression patterns and the activation of certain oncogenes, for instance, c-MYC, RAS and SRC [66, 67].

Taking into account all the above information, one can surmise that the elastic field drives glucose transport through the cell membrane so that it diffuses elsewhere. Thus, one can assume that the amount of biomass transported per unit time from cell i to a neighboring cell *m* is proportional to the difference of elastic potential energy *V*_*i*_ − *V*_*m*_. In addition, the membrane permeability is proportional to the glucose concentration difference, *c*_*i*_ − *c*_*m*_, between the two cells. Then one can write the following kinetic equation [54]:

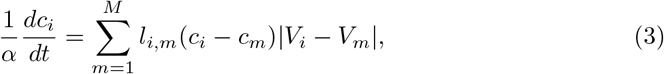

where *l*_*i,m*_ is the contact perimeter between biological cells *i* and *m* and the sum is over all neighboring cells to the *i* − *th* cell.

### Cell Cycle

Recent studies [68] have shown that many genes have a complex oscillation function that depends on the metabolic cycle. In particular, it is known that cell growth can be limited by the amount of glucose in the G1 phase of the cell cycle. Studies in yeast show that there are independent metabolic oscillations in the control of the cell cycle that probably arise from a system of coupled oscillators [69]. In breast cancer, estrogen regulates the expression and function of cyclins, c-Myc, cyclin D1 and cyclin E-Cdk2 which are considered important in the control of G1/S phase progression [70, 71]. These studies suggest that there are molecular coupling mechanisms from metabolism to the cyclin/CDK system that govern the cell cycle. Cyclin E is known to show a periodic expression pattern during the G1 phase of the cell cycle. The levels of this cyclin increase during the late G1 phase and subsequently decrease in the S phase, leading to a significant increase in cyclin D. In the subsequent phases G2 and M of the cell cycle, cyclin D is degraded and cyclin E begins to increase again in the early G1 phase. Other CDK complexes, such as cyclins A and B, undergo periodic patterns opposite to those of cyclin E [71]. These antagonistic processes of the CDK complexes can be explained in terms of non-linear oscillations in which the maxima point to different phases of the cell cycle. Particularly, we could associate one of these maxima with the transitions of the G phases of the cell cycle [72]. Considering that cyclins E and B are the two key players responsible for the out-of-phase oscillations that drive the cell cycle, we can think of the dimensionless Lotka-Volterra system of equations as the mathematical model that describes the cell cycle oscillations.

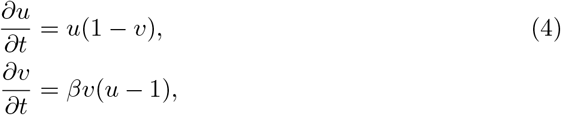

where the functions *u(t)* and *v(t)* represent the concentrations of cyclins E and B, of a cell at time *t*, respectively. The wave shape solution depends only on the local parameter *β* and on the boundary conditions of each cell. This system of equations describes out of phase oscillations with period 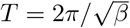.

Considering that each cell has a slightly different period due to fluctuations in glucose local concentration, i.e. *β* = *γc*_*i*_ with *γ* an adjustable parameter. Then, the cell clock cycle of each cell can be estimated by,

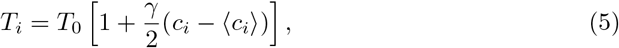

where 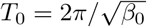is the phase oscillation of basal cells with ⟨*T*_*i*_⟩ = *T*_0_ [73].

The transport of glucose is described by Eq (3) where the elastic potential variations may change locally the amount of glucose in the cells. This fact can significantly change the reproduction cell clock cycle giving rise to an unbalance in the tissue. To take this into account we estimated the time derivative of the glucose fluctuations by taking the derivative of Eq (5).

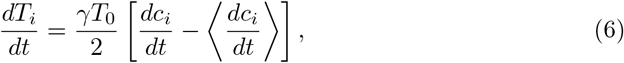

where we consider

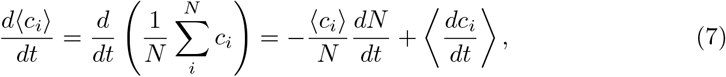

and assumed a homeostatic balance, i.e. *dN/dt* ≈ 0. To estimate the cell clock cycle of each cell we solved Eq (6) coupled to Eq (3).

### Gene regulatory network

Genetic regulatory networks (GRNs) are build based on the biological regulation of complex biomolecular interactions in tissues that consider the role of specialized cells and generally the cellular microenvironment. In the case of pancreatic cancer, a GRN has been identified from the activity of pancreatic cancer cells (PCCs) and pancreatic stellar cells (PSCs), as well as on the interactions between cytokines and growth factors that communicate between them [36]. On the other hand, network motifs in the GRN are important biological structures that describe cascade regulation and feedback between its agents and their behavior can be analysed through Boolean, stochastic, continuous, or delayed dynamics [14–16, 74–76]. In particular, it has been shown that delayed dynamics plays a key role in the description of the dynamics of these genetic networks. It can be used to further understand biological changes as it offers a more complete and detailed view of GRN behavior [36, 76].

We take as a starting point the GRN proposed by D. Plaugher, et. al. [36], where the most relevant genes and molecules in the expression and suppression of signaling pathways related to pancreatic cancer were identified. To understand the underlying mechanisms that arise from hyperplasia or dysplasia we consider a simpler GRN version that contemplates only the stable states related to the apoptosis and proliferation phenotypes of pancreatic stellate cells and cancer cells. We have deleted some interactions and focused only on the birth and death processes that can lead to tissue inflammation. The simplification of the GRN introduced temporal delays in the dynamics of the pancreatic cancer main regulatory genes, however, the stability of the original GRN remains unchanged so that the genetic dynamics of the simplified GRN yields information about the cell cycle in early PanIN 1 and PanIN 2 stages of pancreatic cancer.

The reduced GRN (R-GRN) of pancreatic cancer presented in Figure 2 contains the actions of the cytokines TNFa and TGFb1, which act as regulators and repressors of genes in PSCs and PCCs that culminate in observable phenotypes, such as proliferation, migration, activation, apoptosis and autophagy. In addition, in the apoptosis of cancerous pancreatic cells, the R-GRN considers the caspases (CASPs) as inflammatory response agents and apoptosis processes in cancer cells. Additional, the R-GRN considers as the main genetic agent the RAS gene, since clinical data in pancreatic cancer show that in all PDAC and PanINs cases, this gene is overexpressed as compared to healthy individuals. The R-GRN uses the interactions of the P53 tumor suppressor gene, the p21 gene as CDK inhibitor that acts as checkpoint protein in the G1 cell cycle phase. The EKR extracellular kinases participates in multiple cellular processes such as proliferation, differentiation, and regulation of transcription. Also, the R-GRN contains the BCL-XL gene that regulates cell death and the PIP3 gene that causes translocation of glucose transporters to the plasma membrane and glucose uptake into tissues.

**Fig 1.**
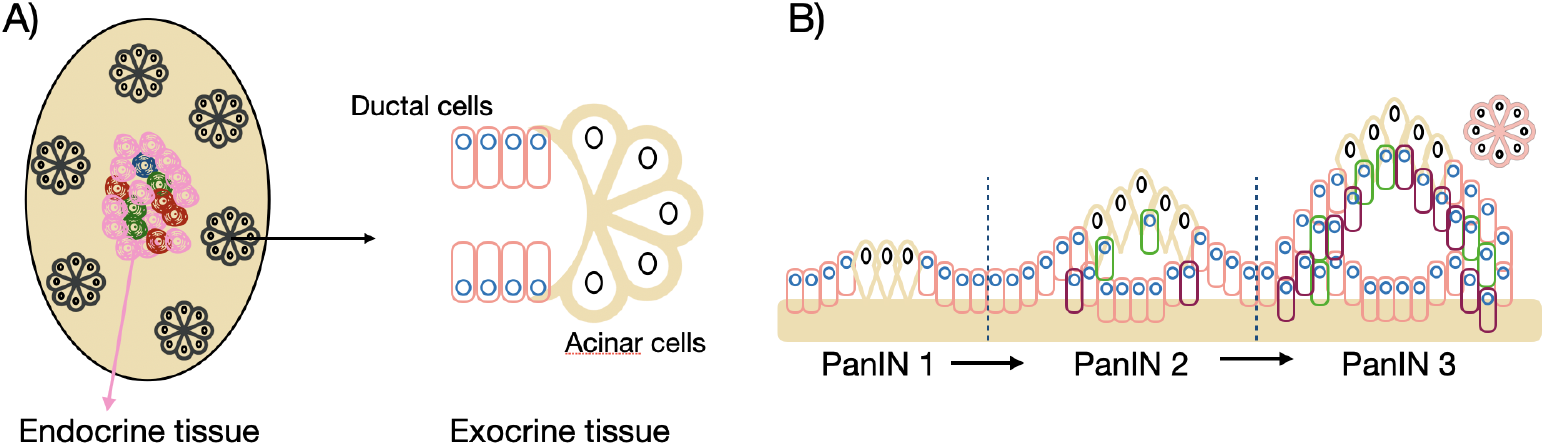
Typical pancreatic tissues. A) Depending on their cell of origin, there are different types of pancreatic cancer. The pancreas can be divided into two tissue types; endocrine and exocrine. Exocrine tissue contains enzyme producing glands, while endocrine tissue contains hormone producing islet cells. In exocrine glands, acinar cells lead to non-ductal neoplasms whereas ductal cells give rise to ductal neoplasms. B) Pancreatic intraepithelial neoplasias (PanIN) and their transitions. The diagrams presented here are inspired in figure 2 of the ref [34].

**Fig 2.**
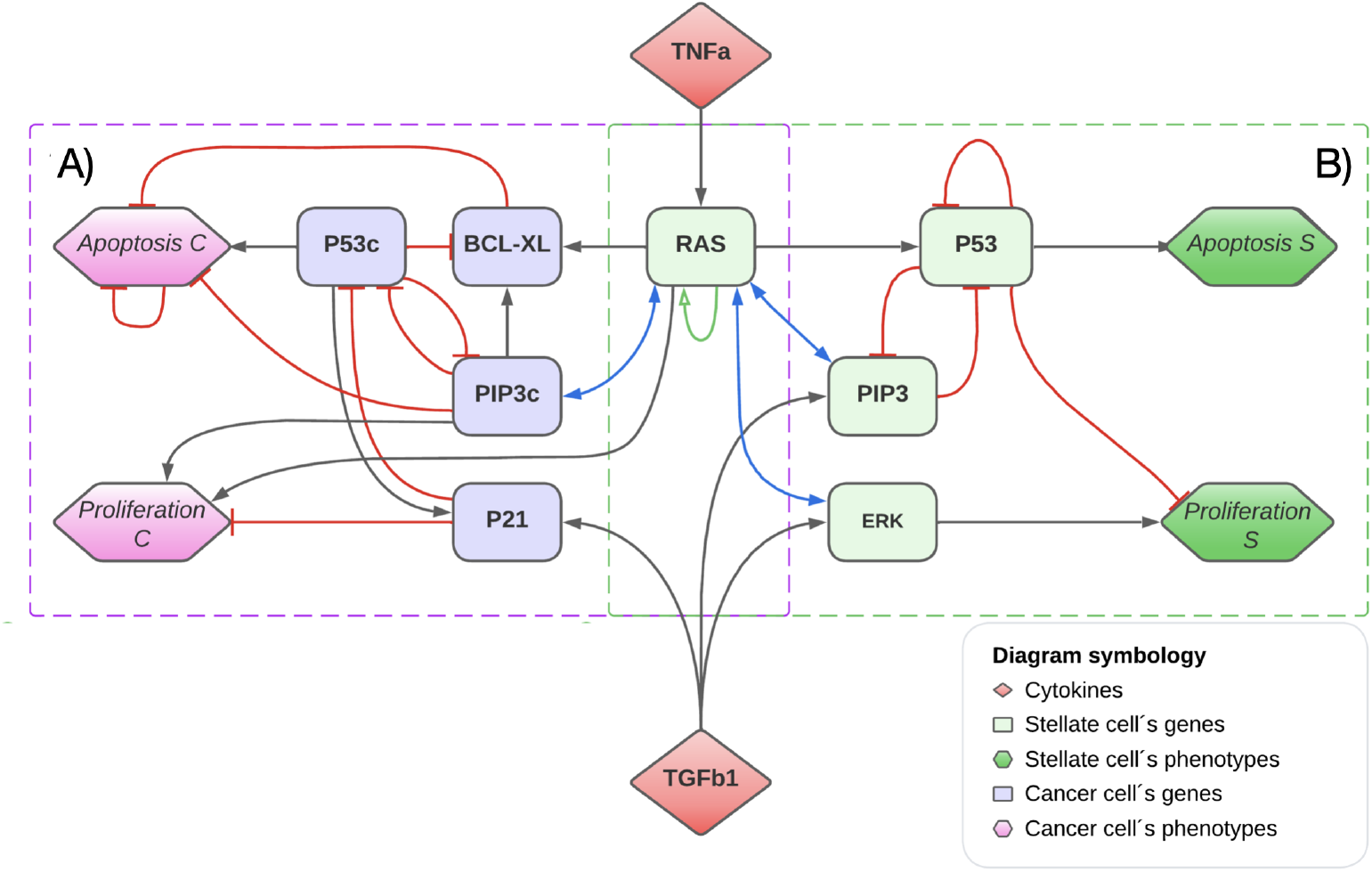
Reduced GRN for pancreas describing the PanIN 1 and PanIN 2 stages. A) On the left side a GRN for pancreatic cancer is represented with purple squares. B) On the right side a GRN for healthy tissues is represented with green squares. The obliques lines represent inhibition actions while the arrows means an activation actions between agents. The letter c at the end of some genes represents the action of a mutated gene.

Note that the KRAS gene is an essential part of the control, as it affects both healthy stellate cells and cancer cells equally. However, the evolution in both cell types are different. Therefore, in order to distinguish the properties of each cell and have a better precision with the experimental evidence, it is convenient to introduce a feedback with time delay that simulates the interaction of the RAS gene with the entire GRN. The previous proposal allows to introduce a free parameter *τ* that will play a crucial role when calibrating the clinical activation times between the different states of the PanIN processes. Thus, the value of *τ* captures the temporal progression of pancreatic cancer, from PanIN 1 to PanIN 3 onset, and provides a better correspondence to apoptosis processes in cancer cells.

### Modeling the R-GRN

Different methodologies and approaches have been developed to model GRN dynamic interactions. For instance, aspects such as cooperation, competition and regulation have been analyzed [36, 77, 78]. Likewise, for the models mathematical formulation, aspects such as simulation of logic gates [79], network motifs [80], design of genetic circuits [81] and modeling of genetic circuits [82] have also been considered. Models based on sets of ordinary differential equations (ODEs) have been proposed to describe complex biological phenomena, such as population growth, enzyme kinetics, among others [83, 84]. Despite being formed by relatively simple systems of ODEs, these models are able to capture many nonlinear aspects of the biological systems. Here we have carefully modified the pancreatic cancer regulatory network studied in reference [36] to obtain an optimized version. In this simplified version of the network we have made sure that the proliferation and apoptosis attractor states remain in the dynamics. To investigate the convergence of the long term temporal dynamics we have analyzed it by using continuous logic gates [85, 86] and have obtained the correct gene expression profiles.

Delayed systems of differential equations have been incorporated to mimic the delayed response of the interactions between the network components [87]. These delays are related to the biological processes of transcription, transduction, and transport of molecules. The inclusion of time-delayed systems of differential equations adds more realism to the model although introduces some difficulties when solving them. Nonetheless, it captures with more precision the temporal dynamics response of the GRN to biochemical reactions and positive feedback circuits between the network components [88].

Logic gates describe the Boolean actions of more than two variables, they regulate several input signals and result in a single output signal. Because of this, they are fundamental elements in the modeling the dynamics of a GRN. In this context, the interactions between genes and proteins in a biological system can be modeled as input and output signals. As a matter of fact, there is a unified Hill-function logic framework, an equivalence between Boolean logic and a superposition of cooperative functions [87]. In what follows we apply this framework to model the actions of the R-GRN. Here we consider a transcription factor *x(t)* that regulates in time the production of a protein *y(t)*. If *x(t)* activates *y(t)*, then the production rate of *y(t)* is proportional to the variations of *x(t)*. The cooperative interplay is modeled in the context of Hill regulation with a parameter n that determines the steepness of the increase of *y(t)* in response to an increase of *x(t)*. To account for the explicit delay in regulation, we replace *x(t)* with *x(t* − *τ*). Therefore, the cooperativeness differential delay equations are:

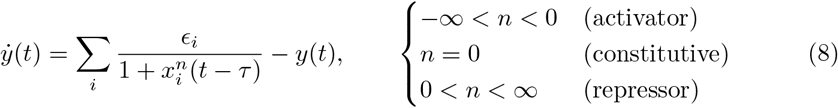

where the sum is over the nearest neighbors of gene *y* in the R-GRN, *ϵ*_*i*_ is the regulation strength, n is the cooperative parameter and *τ* is the time delay. In general, gates can exhibit a high (“on”) or low (“off”) output signal depending on whether the inputs are on or off. Therefore, the GRN parameter space can be divided into two types of logic functions: AND type and OR type. The former functions are activated when both, the x and y regulations are weak, while the latter functions are activated when both regulations are strong. These two types of logic functions can be interchanged at once by applying negation to one or both inputs. The set of delay differential equations that describe the dynamics of the GRN in the Fig. 2 are written in the supporting information. Here, we write down the adimensional equations that are solved (the details about the delays can be found in [79]):

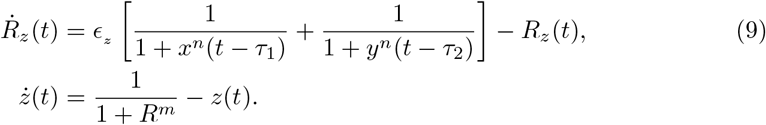

In these equations *R*_*z*_ is the regulation response of variable *z* due to *x* and *y* agents, and *z* is the regulated agent. The R-GRN variables and their representation in terms of Boolean functions is presented in table 1. The activation is represented by OR and the suppression by AND NOT.

**Table 1.**
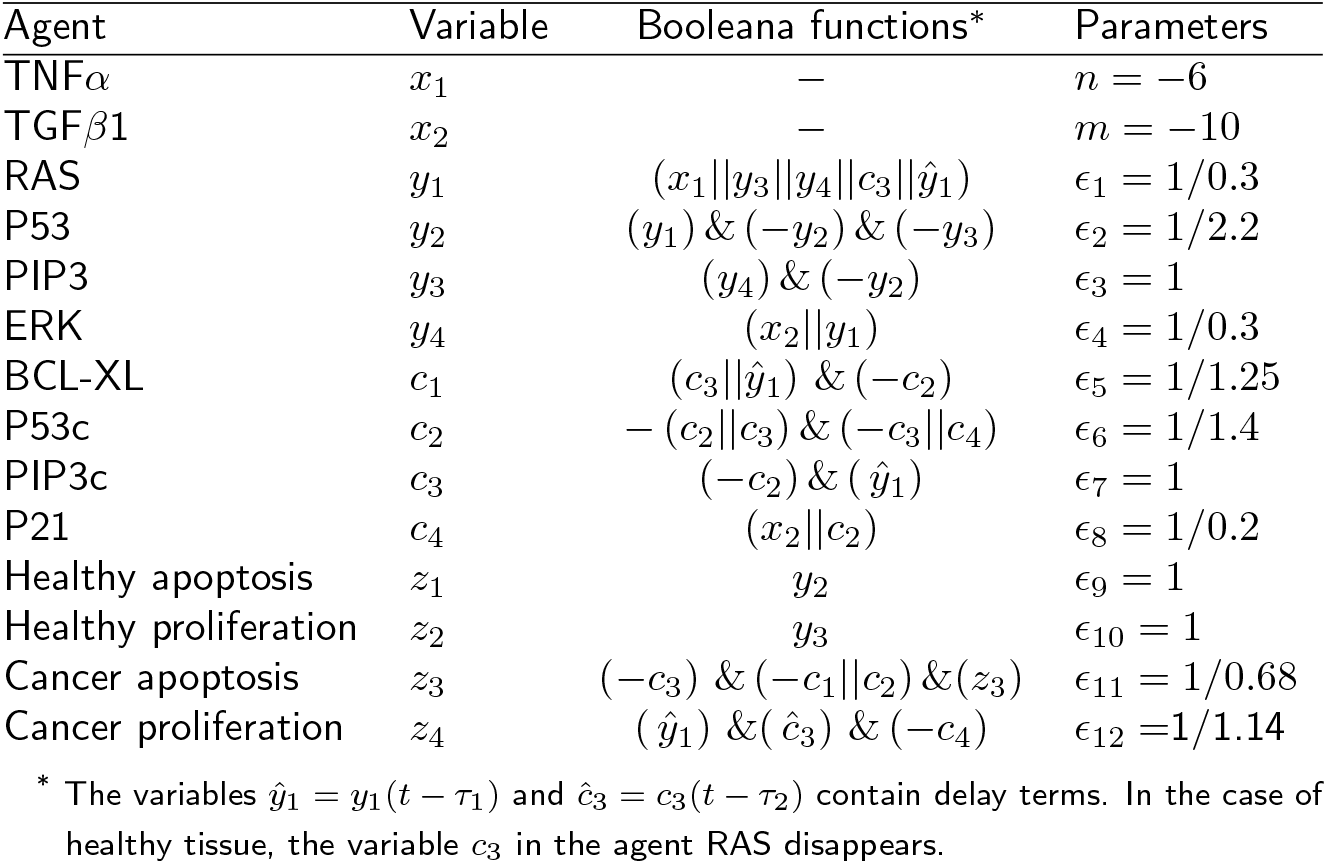
Optimized Boolean functions of cytokines, genes and phenotypes present in the model of the cellular microenvironment in pancreatic cancer.

### Numerical integration

#### Temporal analysis of the PanINs evolution

In a recent study the average residence times at each stage of PanIN were examined [89]. They found that the residence average time in PanIN 1 was 23.7 years, whereas the residence average time in PanIN 2 was 17.5 years. These findings indicate that the transition time from PanIN 1 to PanIN 2 is faster in cancer cells, suggesting that a temporal delay in the GRN dynamics equations is an important point to be considered in the evolution of pancreatic cancer. Because of this, here we have analyzed the GRN dynamics in terms of delay differential equations. In a comparison of the residence times in Ref. [89] between PanIN 1 (*τ*_1_) and PanIN 2 (*τ*_2_) for men (*m*) and women (*w*), one finds that the ratio between both times remains almost constant, that is, *τ*_2,*m*_/*τ*_1,*m*_ ≈ *τ*_2,*w*_/*τ*_1,*w*_ ≈ 0.8. Which somehow shows that the inflammatory nature in PanIN 1 is less progressive and aggressive compared to PanIN 2. From the above, one can assume that cancer cells are responsible for moderating genetic deregulation by generating delays in GRN with a proportion *τ*_2_ = 0.8 × *τ*_1_. Thus, taking into account the mean values reported in the same Ref. [89] for the transitions between PanIN 1 and PanIN 2, we have determined the calibration values of our model as *τ*_1_ = 17.13 and *τ*_2_ = 13.82 years, respectively.

#### Spatial analysis of the PanINs evolution

We have analyzed the genetic-phenotypic expression profiles of PSCs in both, a healthy pancreas and one with the presence of cancer cells (PCCs). Figure 4 shows the GRN phenotypic expressions results for the healthy pancreas. The GRN exhibits a balanced and coordinated genes expression whereas the associated phenotypes, such as proliferation and apoptosis, were at stable and controlled levels. These results indicate that the healthy pancreas is in a state of cellular homeostasis. For the evolution of a healthy pancreatic tissue, we incorporated in the GRN the cytokine cycle since it plays a fundamental role in the genetic regulation and maintenance of homeostasis. By considering a basal cytokines period of *T*_0_ ≈ 36 days [73] and the cytokines concentrations TNFa (*X*_1_) and TGFb1 (*X*_2_) we obtained from the solutions of the Lotka-Volterra equations (Eqs (4)) for cyclins E and B. The Fig 3 shows the activation cycles of cytokines.

**Fig 3.**
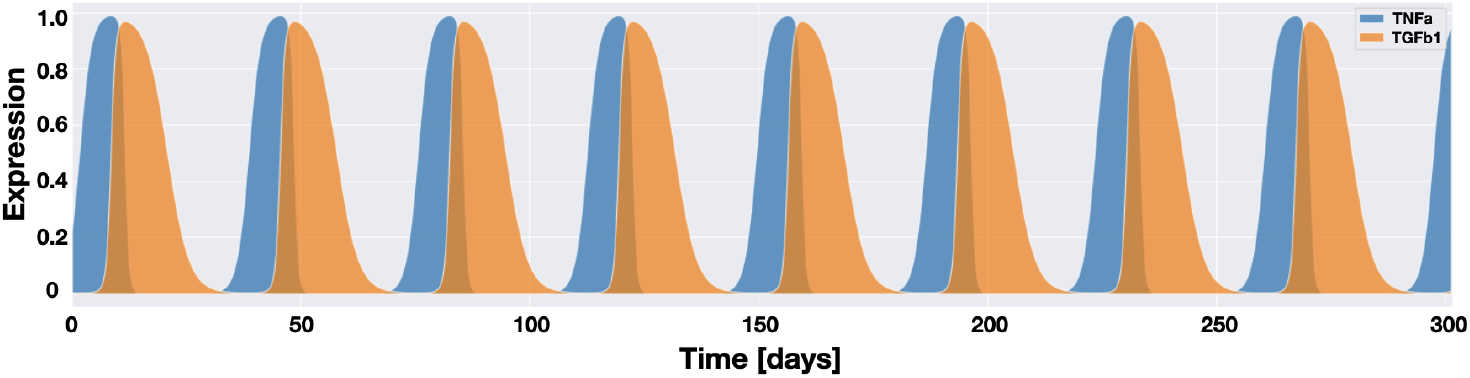
The graphs show the temporal response of the cytokines TNFa (X_1_) and TGFb1 (X_2_) over time. The regulated action of cytokines can be an indicator of homeostasis processes.

#### Spatial relaxation

To make an easy quantitative description of the inflammation of the acinar-ductal system here we used a Voronoi diagram construction to physically represent this system. Taking into account that pancreatic acinar cell injury triggers the synthesis and release of pro-inflammatory cytokines and chemokines [90–94]. The values of the parameters *K*_*c*_, *K*_*v*_, *k, α* and *γ* where taking to illustrated the dynamical behavior of the system. Considering that the parameters *K*_*c*_ and K_*v*_ are related to the elastic modulus E of the cells [54] and with the healthy length of the acinar-ductal cells, we choose *K*_*c*_ = 0.3 Pa-m, *K*_*v*_ = 0.06 Pa/m, due to elastic modulus of living cells is of the order of *E* ∼ 5–10 KPa, as reported in ref [95] and the diameter of the acinar-ductal cells varies between 10–24μm [96]. The friction parameter of Eqs (2) is choosen as *k* = 0.001 Pa-m-s and *γ* = 60 corresponds to one basal oscillation *T*_0_, when the glucose mean concentration is ⟨c_*i*_⟩ = 0.5.

We introduce a quantification of the inflammation in the acinar-ductal system and two key biophysical parameters: the cell size domain represented by its area *A*_*T*_ and the basal cell size represented by its Voronoi area, *Â*_*cell*_. Accordingly, the inflammation was quantified by measuring the ratio, *A*_*T*_ /*Â*_*cell*_, between these two areas. Thus, when the geometric domain occupied by the cells is under pressure due to inflammation, (*A*_*T*_ /*Â*_*cell*_ > 1). On the contrary, when the system tissue is relaxed and inflammation is absent, cells have enough space to move around and migrate, so that, (*A*_*T*_ /*Â*_*cell*_ < 1). Cell proliferation increases the cell density and therefore contributes to the growth of the duct in the acinar-ductal system which in turn reduces the inflammation and maintains an adequate balance in the tissue. In this situation, the following condition should be satisfied *A*_*T*_ /*Â*_*cell*_ > *A*_*up*_, where *A*_*up*_ is a threshold value that determines the duct elongation. In the supporting information section we present the calculations that lead to the duct’s length.

#### Spatial cell distribution

Cell proliferation is a fundamental process in the growth and development of biological tissues. It determines the shape, density and size of the growing tissue and plays a crucial role in the formation and maintenance of the structure and function of organs. Because of this cell division process should be considered in a tumor growth model. In doing so we have considered the following four steps: *i) identification of cells that proliferate*; *ii) generation of the corresponding daughter cells*; *iii) Mechanical relaxation and adjustment of the tissue as a result of its growth*, and *iv) Cell death*. In what follows we describe each of these four steps.

##### i) Cell proliferation

According to Eq (6) those cells that have a clock cell cycle larger are those that have a local glucose concentration, *c*_*i*_ larger than the average glucose concentration < *c*_*i*_ >. Taking this into account we identify the cells that proliferate as those in which their internal clock cell cycle (*T*_*i*_) is greater than the basal clock cell cycle (*T*_0_). That is, cell division happens when 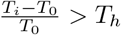. Here *T*_*h*_ a threshold value that is chosen such that the area of the mother and daughter cells adjust easily to the tissue mechanical equilibrium. We found that the threshold optimal value was *T*_*h*_ = 0.15. This threshold has been kept fixed in all numerical simulations.

##### ii) Generation of daughter cells

Cell division proceeds through a precisely timed and carefully regulated stages of growth, DNA replication, and division that produce two genetically identical cells. Here we have considered a geometrical procedure that leads to the cells descendants. Let us assume that the shape of the two-dimensional mother cell is circular with area *A*_*i*_, so that its radius is 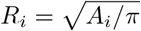. In order to guarantee that the area of each daughter cell is one half of the original mother cell area, *A*_*i*_, the centers of the daughter cells should be distant by R_*i*_/2 from the center of the mother cell. Then, each daughter cell will have a radius 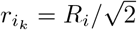, with *k* = 1, 2, and their initial positions are chosen in a random direction. In addition, we shall assume that the kinetic energy, *E*_*c*_, of the mother cell is shared with the daughters cells. After division the masses of the daughter cells are one half of their mother cell, that is, *m*_1_ = *m*/2 and *m*_2_ = m/2. To ensure that the daughter cells have an adequate amount of energy for their subsequent development and motility we assume conservation of momentum and kinetic energy during the division process. Then the magnitude of the velocity of each daughter cell becomes 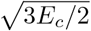. Additionally, we assume that during cell division the glucose concentration is shared by one half each daughter cell.

##### iii) Relaxation and adjustment

After cell division occurs, the tissue relaxes generating elastic interactions with the daughters’ neighboring cells and their surroundings until an equilibrium mechanical configuration in the tissue is reached. To have synchrony between the mechanical equilibrium and the basal cell reproduction period T_0_ we shall assume that the mechanical relaxation process takes the same time T_0_.

##### iv) Cell death

Cell death is a physiological process to ensure correct development or tissue homeostasis, nonetheless, it can be considered as a pathogenic mechanism that undermines normal organ function and leads to local or systemic inflammation. Cell death can be classified into two large categories: accidental cell death and regulated cell death [97, 98]. Regulated cell death plays a dual role in pancreatic cancer and has been shown to have both pro-tumorigenic and tumour-suppressive effects. At the molecular level in PDAC various oncogenes or tumour suppressor signals determine the sensitivity of cell death modalities. The main types of regulated cell death that have been identified in PDAC are: apoptosis, necroptosis, ferroptosis, pyroptosis and alkaliptosis [99]. Since the detailed mechanisms of cell death in PDAC are still to be discovered here we simulate the cells death process by considering their internal cell cycle clock. To this end we implemented the following steps: *a) Identification of candidate cells for cell death*. The candidate cells for death are those that have an important difference in their internal cell cycle clock T_*i*_ as compared the basal clock cell cyclel, T_0_; that is, when 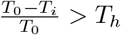. When this inequality holds the cell i is eliminated from the tissue. The previous ratio inequality is an indicative that cells in latent states will die because their internal clock cell cycle is shorter than the normal one. Voronoi points associated with these cells are eliminated. *b) Cell elimination*. After cell removal, the Voronoi diagram is locally recalculated to maintain a valid and stable configuration of neighboring cells, simulating the process of local tissue restitution.

The normal evolution of the system begins by allowing the action of mechanical and elastic fields in the tissue. Subsequently, transport of glucose by diffusion gives rise to a gradient in the system. By solving the transport Eq (3) we obtained the normalized glucose concentration. Next, the internal clock of the cells advances by the amount *Δt*, an increment of the simulation time, that is, 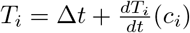, until a time of *T*_0_/2 is reached. When this happens, the internal clock cell cycle of each cell is checked out and a determination is made if this cell either divides or dies. Once the cells fate is determined an iteration of Eqs (2) is carried out in order to homogenize and stabilize the system. During this process, cells interact with each other and with their environment, adjusting their elastic equilibrium position to minimize the potential energy and reach a tissue stationary state.

## Results

### Emergence of phenotypes from GRN dynamics

In Fig 4A it is observed that the gene RAS expression level remains high whereas the gene P53 expression level is low, which contributes to maintain a balance in stellate cell apoptosis (Apop S) and proliferation (Prolif S). The gene PIP3, that regulates cell proliferation maintains its expression at moderate levels, preventing excessive proliferation of pancreatic stellate cells. In addition, gene ERK shows pronounced activation values suggesting a greater response to the growth signals. In Fig 4B it is shown that the proliferation phenotypic expression reaches a peak as pancreatic stellate cells experience an increase in their proliferation rate. This is an indication that cell growth as well as renewal of the pancreatic microenvironment are underway. Alternatively, the phenotypic expression of apoptosis reaches a peak suggesting a stable high rate of programmed cell death. This behavior yields a bistable cycle between proliferation and death which is crucial for the dynamic balance in a healthy pancreas. In this way, healthy tissue cells respond and adapt to changes in their environment in a controlled manner, preventing the development of cellular alterations. This is a relevant feature in the GRN dynamics of a healthy pancreas. A different type of dynamics breaks the homeostatic balance and can lead to the development of inflammation and/or stress processes.

**Fig 4.**
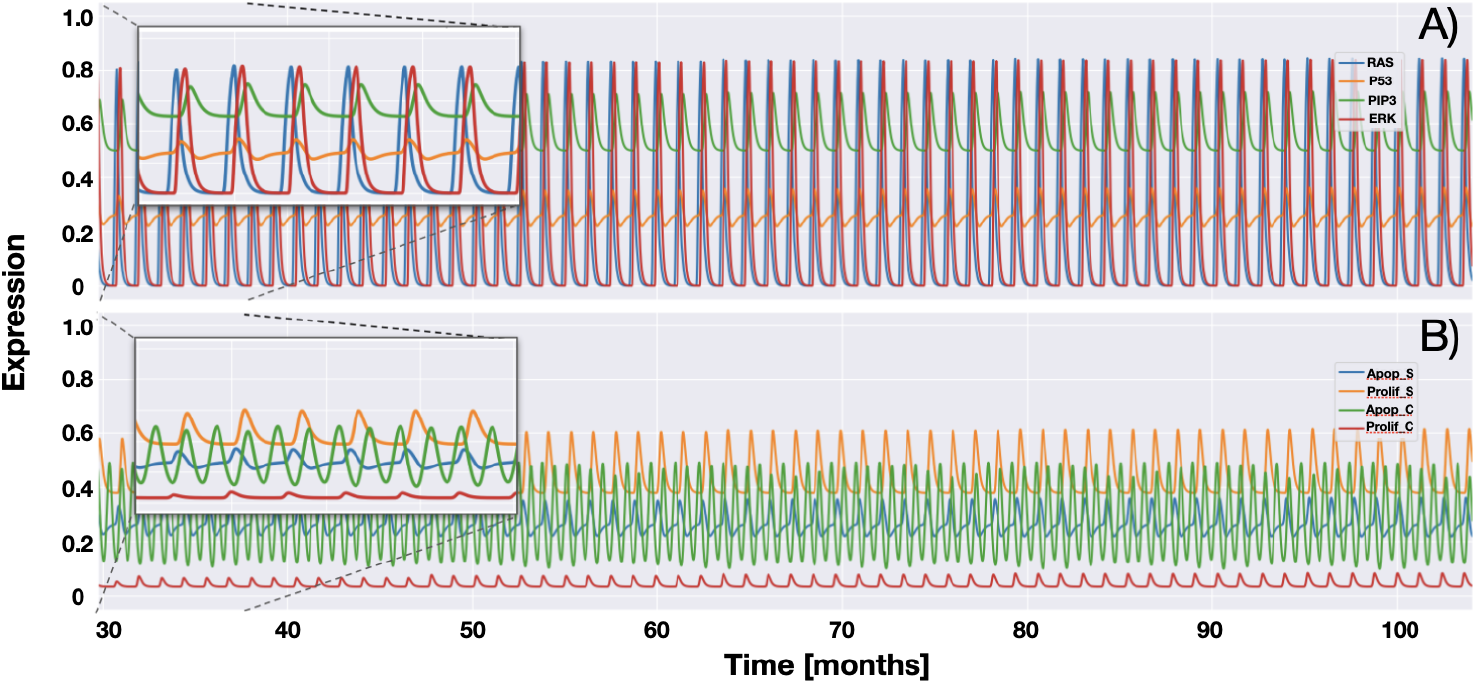
Genetic-phenotypic expression profile for genes in the GRN of an isolated healthy pancreas. A) The oscillation behavior of the genes RAS, P53, PIP3 and ERK. B)The states of cell phenotype due to proliferation (Prolif S) and apoptosis (Apop S) of healthy cells, and proliferation (Prolif C) and apoptosis (Apop C) of cancer cells.

Fig 5 shows the genetic-phenotypic expression profiles for the GRN genes once a pancreatic cancer has developed. Figs 5A and 5B show an increase in the genetic-phenotypic expression profile of the PSC and PCC cells, respectively. These results indicate that the introduction of the delayed times *τ*_1_ and *τ*_2_ in the GRN kinetic equations is critical for the description of the pancreatic cancer progression. As a matter of fact, the delayed time *τ*_1_ marks the transition from the PanIN 1 to PanIN 2 states, while *τ*_2_ marks the change from the PanIN 2 state to an advanced cancer PanIN 3 state. For PanIN 1 state, (0 < *t* ≤ *τ*_1_), one observes that the GRN maintains a profile similar to that of a healthy pancreas in the PSCs cells. However, for the PCC cells, there are some incipient alterations in some genetic agents. For instance, cancer cell proliferation (Prolif C) increases whereas cancer cell apoptosis (Apop C) decreases, which are two hallmarks of cancer development. The evolution to the PanIN2 stage (*τ*_1_ < *t* ≤ *τ*_2_) yields the activation of the ERK gene, which in turn leads to an increase in the activity of the cell growth pathway. In Fig. 5C one observes that as time approaches the value (*τ*_2_) the proliferation of cancer cells (Prolif C) oscillates with larger amplitude and higher frequency. This suggests that there is a stronger and sustained activation of cancer cells, while PSC cells seem to maintain a relatively stable behavior. It was observed that in the PanIN 1 stage, cancer cells show a slow increase in proliferation and a decrease in apoptosis. This suggests that cancer cells appear to be in a dormant state since they proliferate at slow rate. On the other hand, in the PanIN 2 stage the oscillatory behavior in the cancer cells proliferation suggests a transition to a higher proliferation rate, which may lead to the onset of localized tumors in the pancreas [100]. Furthermore, significant deregulation of the ERK gene could indirectly contribute to increase its mutations by increasing proliferation. This deregulation deactivates the apoptosis mechanisms in cancer cells, which favors their survival over healthy cells and leads to tumor growth (see the inside figures).

**Fig 5.**
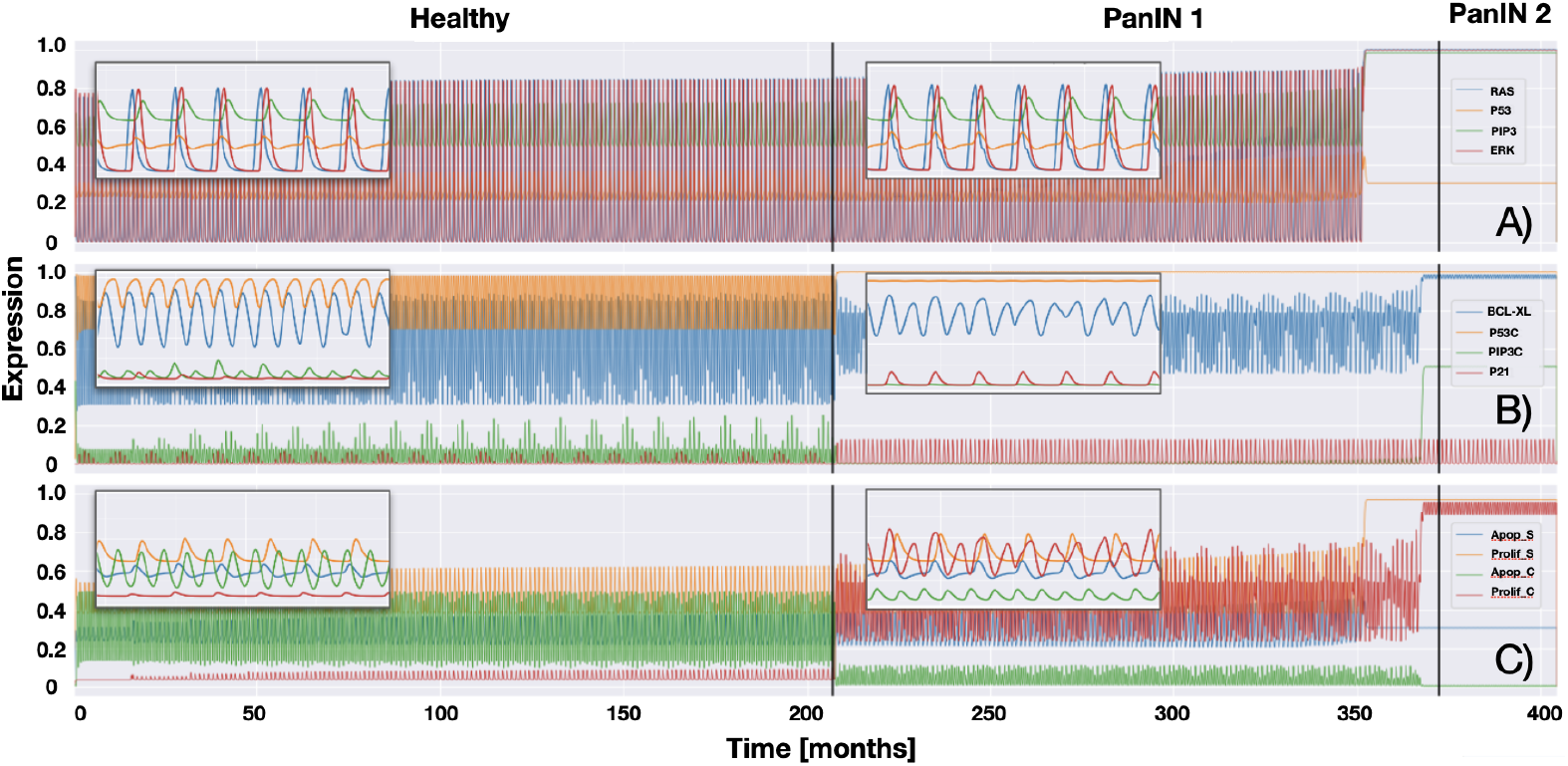
Genetic-phenotypic expression profile for the GRN genes of a pancreas with cancer. The genes are in two states: activated or inactivated. A) The oscillation behavior of the healthy genes RAS, P53, PIP3 and ERK. B) The oscillation behavior of the cancer genes BCL-XL, P53C, PIP3C and P21. C) The states of cell phenotype due to proliferation (Prolif S) and apoptosis (Apop S) of healthy cells, and proliferation (Prolif C) and apoptosis (Apop C) of cancer cells

Fig 6 shows the distribution and evolution of cells, without considering cell division and death, in the tissue. They are subjected to mechanical and plastic forces such as cell adhesion and edge tension. Fig 6A represents an initial state where cells are randomly distributed. The mechanical and plastic forces acting on cells generate deformations and changes in their shape and positions that yield mechanical stress in the tissue until an equilibrium state is reached. During the relaxation process the glucose concentrations in the cells tend to distribute evenly throughout the tissue. While this mechanical relaxation process happens, GLUTs facilitate the diffusion of glucose across cell membranes. The glucose concentration distribution is shown in Fig 6 and can be quantified by looking at the color bar located on the right side of the figure. Notice that stressed cells show higher levels of glucose concentration. By construction of the Voronoi diagrams cells are in contact giving rise to glucose concentration gradients in the tissue which ensures that cells have an adequate glucose supply for their metabolic processes and growth.

**Fig 6.**
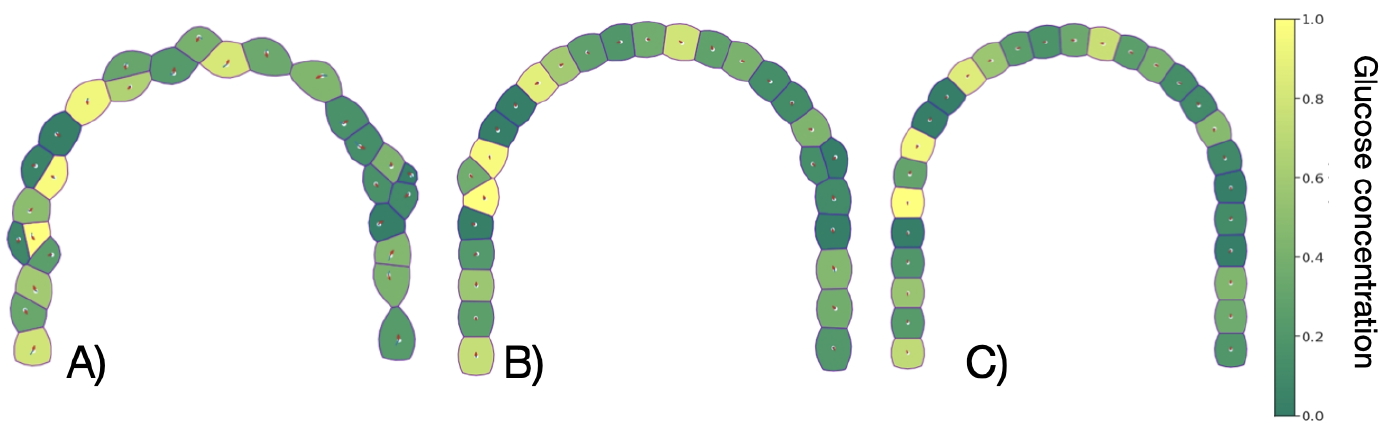
Distribution and evolution of cells subjected to their own mechanical and plastic forces without considering cell division and death. A) Initial state where cells are randomly distributed. B) Intermediate state. C) Equilibrium state. Glucose concentration gradients are also shown and can be quantified with the color bar located on the right side.

### Healthy pancreas

A healthy pancreas is free of inflammation and has a balanced distribution of gene expression and cellular phenotypes. The basal cells cycle is synchronized at the value T_0_ = 36 hours [73]. This means that there should be a compensation between the processes of cell division and death that maintains homeostasis in the system. Recent findings indicate that pancreatic stellate cells (PSCs) play an important role in normal pancreas function, as well as in response to disease and damage. In the healthy pancreas of an adult, PSCs are present in small amounts, nonetheless, they are ‘quiescent’ and regulate extracellular matrix (ECM) production. On the other hand, activated PSCs proliferate rapidly and undergo substantial changes in gene expression. They are perhaps the most abundant cell type of a damaged pancreas. Because of this PSCs have been extensively studied, mainly for their role in pancreatitis and cancer [101, 102]. In cancer, cell-cycle entry is constantly driven by deregulated mitogenic signals, that results in aberrant proliferation. Recent studies in pancreatic cancer cells have found several gene mutations that influence cell-cycle entry [103, 104].

The detailed mechanisms, that govern the process of PSCs phenotype transition in the distinct phases of pancreatic inflammation and tumorigenesis, are a little understood. The delay introduced in the GRN kinetics emulates the KRAS, CDKN2A, TP53, and SMAD4, genetic mutations assuming that each mutations introduces a desregulation in the GRN performance. Usually, KRAS mutations are observed in early lesions (PanIN-1) followed by CDKN2A mutations (PanIN-2), whereas TP53 and SMAD4 mutations (PanIN-3) drive tumorigenesis further, which lead to advanced pancreatic cancer.

### PanIN 1 (Low-grade dysplasia)

Pancreatic cancer precursor lesions at PanIN-1 stage are characterized by the onset of inflammation process during which healthy cells are transformed into precancerous cells. Because of this we have considered different inflammation scenarios based on gene expression and on cell proliferation to emulate the effects of inflammation. The acinar/ductal system evolution through the PanIN-1 stage starting from a healthy state is shown in Fig 7. Fig 7A shows a healthy acinar/ductal system followed by light and moderate inflammation states that are shown in Fig 7B and Fig 7C, respectively. There one can observe that as a result of inflammation, cell proliferation and the cellular ratio area/volume increase. A typical process that characterizes the PanIN-1 stage is that during duct elongation, the increase in the cell area/volume contributes to relieve stress on pancreatic cells. In the figures it is also observed that during the inflammation process glucose diffuses through some regions of the acinar-ductal system generating glucose concentration gradients. There are regions where glucose concentration diminishes significantly pointing to cellular high consumption of glucose. These results indicate that changes in cellular metabolism have ocurred as a result of inflammation.

**Fig 7.**
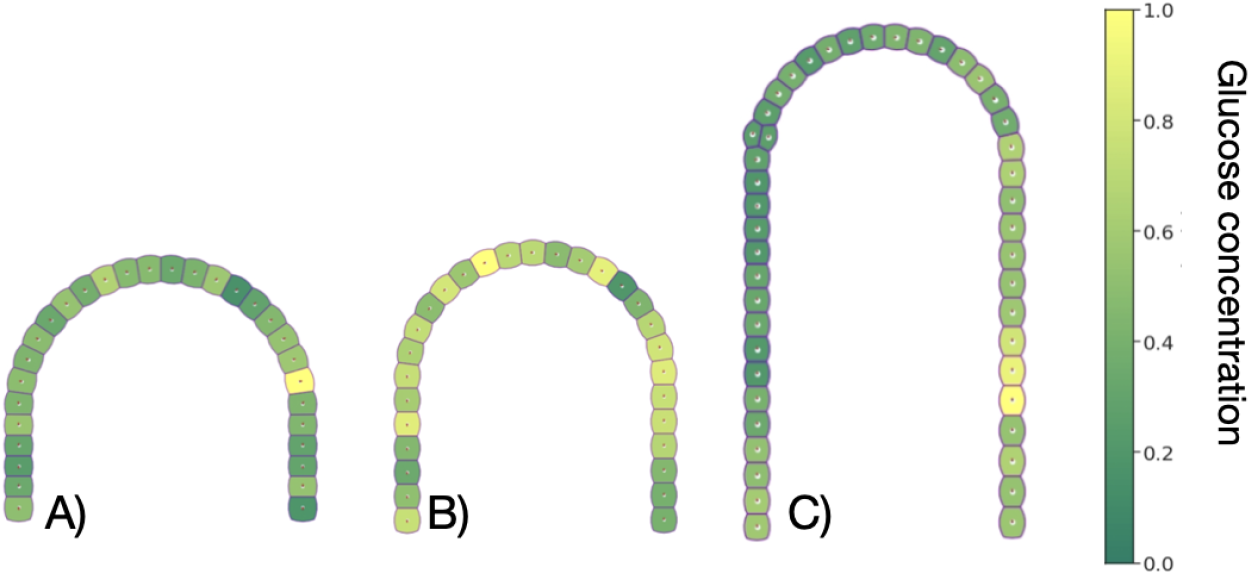
Evolution of an acinar-ductal system: A) healthy B) soft inflammation and C) moderate inflammation. Glucose gradient concentration is indicated in the figures according to the color bar on the right side.

### PanIN 2 (Moderate-grade dysplasia)

PanIN 2 lesions appear to be columnar and contain nuclear changes within the cells such as the loss of nuclear polarity, nuclear crowding, different nuclear sizes, nuclear hyperchromasia and nuclear pseudostratification [105, 106]. During the progression from PanIN 1 to PanIN 2, mutations in CDKN2A gene occur frequently which are associated with the loss of p16 expression [107]. As a result, progressive increase in cell proliferation and survival of cancer cells is observed during the transition from normal ducts to PDAC [108]. Because of this, in the present quantitative model at least one cancer cell has been added to the cellular dynamics in the evolution from PanIN 1 to PanIN 2. These cells are able to proliferate since they are no longer in a quiescent state. To understand how the density of cancer cells affects the proliferative activity of the system as well as the rate of progression of PanIN 2 lesions we have analyzed the response of the system to the inclusion of cancer cells in four representative biological scenarios. Fig 8, shows four years cancer evolution structures, panels A) - D), initiated with six, ten, fourteen and eighteen cancer cells, respectively. The structures were initiated in healthy tissue with no inflammation at all. The structure presented in Fig 8A suggests that the six initial cancer cells remain in a quiescent state because cancer cell proliferation rate is negligible. In Fig 8B it is shown a structure that was initiated with ten cancer cells. It is observed that cancer cell proliferation originates at different places indicating that cancer cells are no longer in a quiescent state. Fig 8C shows a structure originated with fourteen cancer cells. In this case we observe that after four years, a moderate density of cancer cells triggers a significant tumor growth. In fact, we can realize that there occurs a local formation of small tumors. The structure shown in Fig 8D initiated with eighteen initial cancer cells, after four years of progression, leads to a high proliferation rate that triggers the rate of tumor growth. The inflammation index increases to severe levels and triggers the transition to PanIN 3. In this stage there should be a greater degree of aggressiveness as compared to the structures shown in panels A) - C). As expected, these results indicate that as the initial density of cancer cells increases there will occur the formation of more localized tumors with greater density that eventually yield a large and dense tumor.

**Fig 8.**
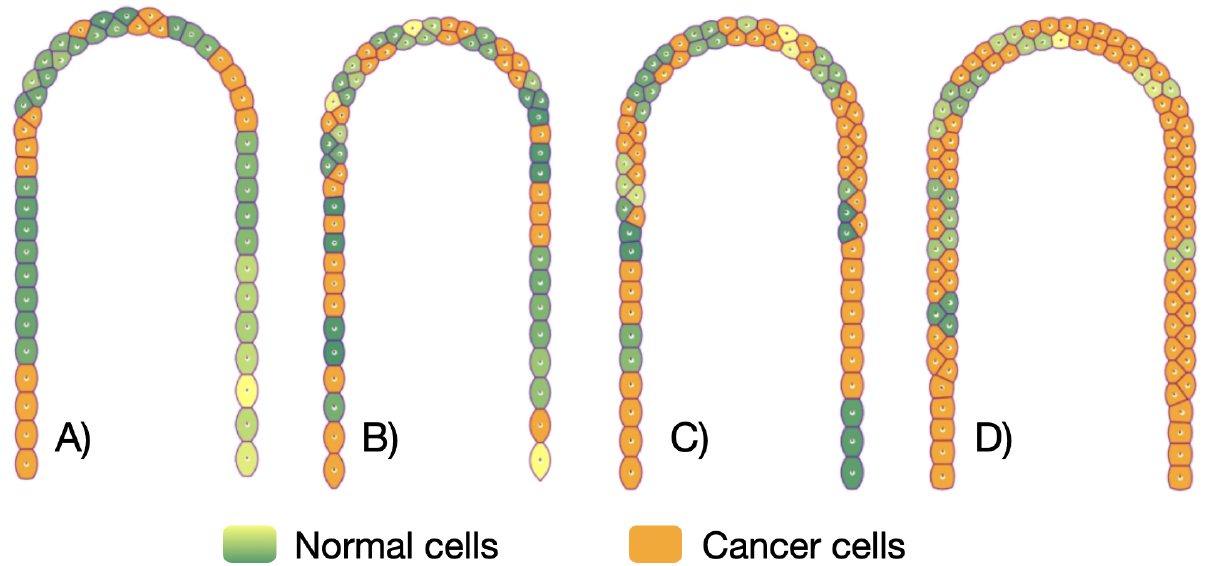
Four years cancer evolution structures initiated with A) six, B) ten, C) fourteen and D) eighteen cancer cells. It is observed that inflammation evolves from moderate in panels (A) and (B) to severe in panels (C) and (D).

### PanIN 3 (high-grade dysplasia)

These lesions rarely are flat, their structure is micro-papillary or papillary and are the result of neoplastic epithelial cell proliferation. They form large nests perforated by many rounded different-sized spaces that eventually give rise to small clusters of epithelial cells into the lumen. Cytologically, these lesions are characterized by a loss of nuclear polarity, dystrophic goblet cells (goblet cells with nuclei oriented toward the lumen and mucinous cytoplasm oriented toward the basement membrane), mitoses that may occasionally be abnormal, nuclear irregularities, and prominent (macro) nucleoli. The lesions resemble carcinoma at the cytonuclear level, but invasion through the basement membrane is absent PanIN 3 lesions have a complex structure with a papillary morphology and enlarged nuclei [109, 110]. They may form clusters of cells that are removed from the epithelium into the lumen of the duct [106].

In addition, it has been found that many cellular signals and pathways contribute to the activation of PSCs, for instance, TGF-*β*, platelet derived growth factor (PDGF), MAPK, Smads, nuclear factor κB, to mention only a few [111].

## Discussion and conclusions

The model presented and analyzed here can mimic a healthy pancreas with a homeostatic state and a balanced cellular dynamics. The cells in the organ proliferate in a controlled manner and respond to programmed apoptosis orderly and in a sequential manner. Furthermore, the model showed that no significant changes occur in the cellular structure at the macroscopic level and that glucose concentrations remain relatively stable. The latter suggests that relaxation of the system is a process in which cells adjust their cell cycle to synchronize it with their neighbors and return to equilibrium. This cellular synchronization could be considered as a form of cellular cooperation, where cells work together as a whole to maintain a healthy state and prevent uncontrolled proliferation. This process occurs through a complex network of interactions and factors so that when there happens a failure it may affect its coherence and stability. For instance, a network failure may lead to stress and inflammation of the tissue which may perturb glucose sources, lead to hypoxia among other factors.

During the growth of the acinus in response to inflammation in the early PanIN 1 stages, the model showed that elongation of the acinar duct and changes in the spatial distribution of cells lead to a stress reduction. Furthermore, to cope with the stress caused by inflammation, an increase in cell proliferation and growth of the acinus structure was observed. We can consider this phenomenon as a resilience effect of the system in PanIN 1, since the acinar-ductal system has the capacity to recover from an inflammatory state and return to a state without inflammation. Likewise, when the system is allowed to relax and undergo slight growth in the acinus (especially in the case of moderate inflammation), the pancreas can return to its stable, healthy state without resorting to cell proliferation.

The presence of a small number of active cancer cells due to the accumulation of genetic alterations (PanIN 2) is sufficient for tumor formation. We observed that the proliferation rate of cancer cells increases as the proportion of these malignant cells grows in the system. Additionally, cancer cells compete for available space to grow as local tumors while healthy cells in the pancreas act as barriers that delineate one tumor from another. Additionally, the model shows that progression of cancer cells is sensitive to the initial proportion of these cancer cells in the system. When a greater number of cancer cells are present in the stages PanIN 1 and PanIN 2, the rate of progression of PanINs to more advanced stages is accelerated. This is interesting as it suggests that controlling cancer cells in early stages (PanIN 1) can prevent rapid progression of pancreatic cancer. An increase in the proportion of cancer cells in the pancreas is an indication of the transition to PanIN 3. In this advanced stage, inflammation increases to severe levels suggesting the possibility of tumor hypoxia and, if boundaries are breached, points to a greater potential for invasion throughout the pancreas acinar-ductal system. It should be noted that spatial dynamics shows emergent behavior as cancer cells interact and compete for the available space to grow and divide. This cellular competition has important implications in the formation and expansion of tumors that proliferate depending on the aggressiveness and proportion of cancer cells while healthy cells act jointly and synchronized to slow growth by acting as natural barriers.

On the other hand, gene expression profiles show a snapshot of the transformation from a healthy pancreas to a pancreas with cancer, revealing the relevance of changes in gene expression in the progression of this disease. Our findings presented in Fig. 5A underline the relevance of several genes in the GRN that undergo notable alterations throughout different stages of pancreatic cancer progression. In particular, we have identified KRAS/RAS as a critical player in the initiation and growth of pancreatic cancer. The deregulated activation of this gene that drives cancer cell proliferation emerges as an essential starting point in the early stages of the disease. Furthermore, variation in P53 expression sheds light on its contribution to the evasion of cellular controls and contributes to cancer cells survival at later stages. In Fig. 5B, deregulation of PIP3 at early stages, especially in PanIN 1 stage, offers important clues about its involvement in the inhibition of pro-apoptotic signals, which promotes cell survival and facilitates the transition to more aggressive states. At the PanIN 2 stage, ERK overexpression emerges as a crucial signal, with the potential to activate the RAS pathway and promote cell proliferation and malignant transformation.

Activation of PIP3 and gradual inactivation of CASP in early stages may facilitate more effective cell survival and proliferation, paving the way to the transition to the more aggressive PanIN 3 stage. At PanIN 3 stage, overexpression of RAS, together with permanent inactivation of CASP, can trigger evasion of cellular control and apoptosis, driving tumor growth and cancer progression. These discoveries not only enrich our understanding of pancreatic cancer progression, but also present exciting opportunities for early diagnosis and effective treatment. Prominent genes such as KRAS/RAS, P53, PIP3 and ERK emerge as potentially useful biomarkers for monitoring the disease evolution, given their correlation with the phenotypic changes observed at the different stages.

The proposed model has the potential to be extended by considering the immune system against the transformation of healthy cells to cancer cells through a better comprehension of the regulatory networks that describe inflammatory processes. Our objective has been to show that the spatio-temporal evolution of cells in the pancreas has its bases in the inflammatory system. Therefore, a better understanding of this dynamics may improve the clinical practice by providing valuable information for the early diagnosis as well as the development of personalized therapies.

## Supporting information

### Differential equations for GRN dynamics

The following set of differential equations describes the dynamics of the GRN presented in Fig 2. It is important to note that this logic scheme is an approximation for functions and the addition of two Hill terms provides a form of fuzzy logic where the values of ϵ close to 1 simulate inputs close to 1 and values ϵ close to 0 create null response. Therefore, given the stability of the GRN in its Boolean form, it is decisive to choose a set of values of ϵ and preserve the dynamic stability of the GRN [79]

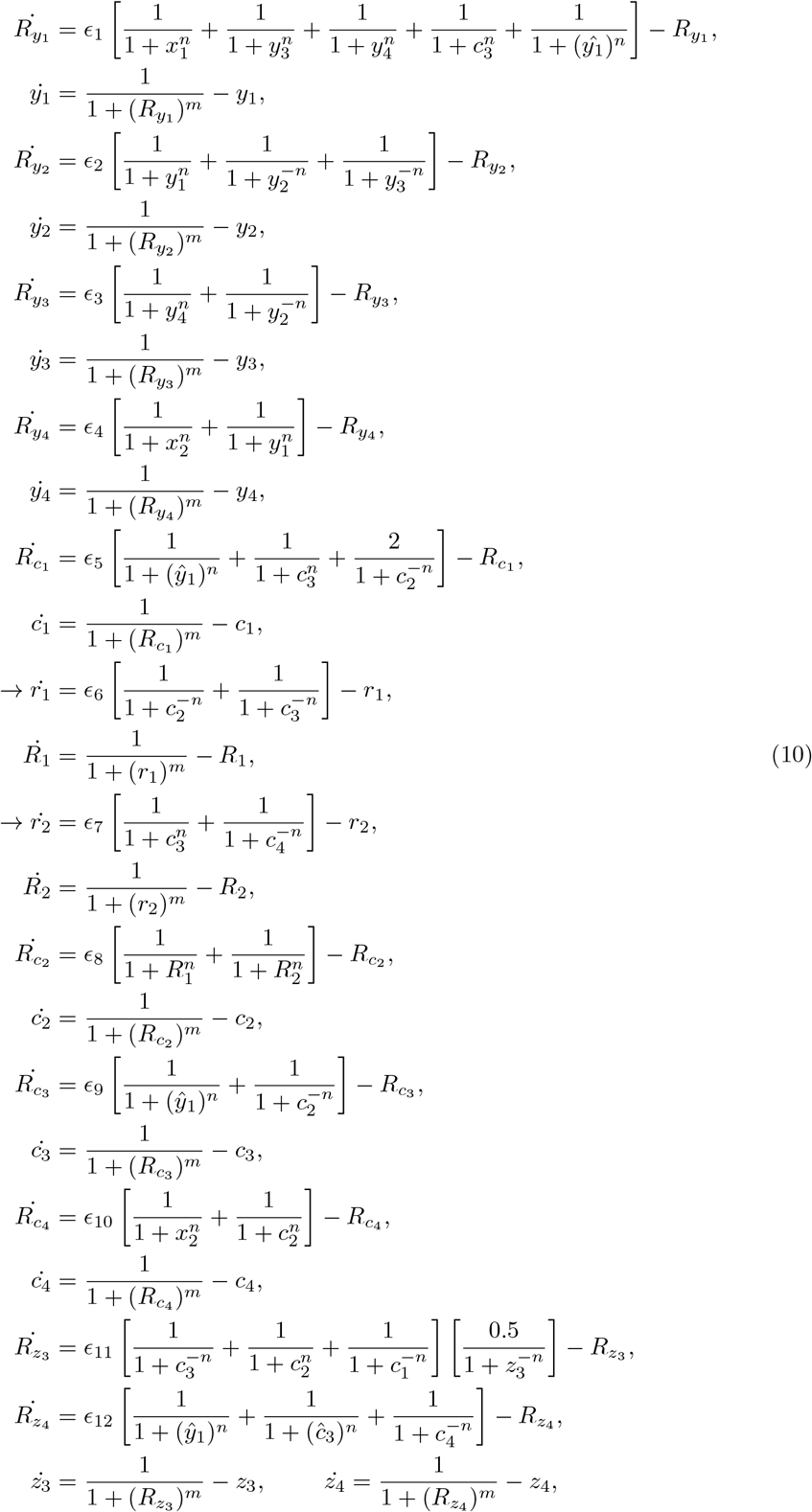

where r_1_ and r_2_ are auxiliary functions that are used to obtain the state of agent P53c.

### Geometric domain of epithelial cells in the acinar-ductal region

The acinar cell structure can be visualized as a *U*-shaped structure that we can mathematically define in terms of some vector functions 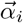 of the form:

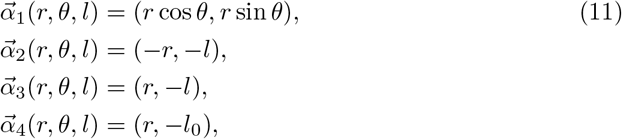

where *r* is the radio defined in *r* ∈ {*r*_*min*_, *r*_*max*_}, and *r*_*min*_ = 2, *r*_*max*_ = 3 are the minimum and maximum radii of the circular shape, θ ∈ (0, π) and the ductal length is l ∈ (0, l_0_) where l_0_ ∈ [1, ∞) represents the vertical shape. The parameter l is a model degree of freedom that allows us to estimate the inflammation of the system. The function 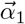 represents a semicircular curve that symbolizes the curved part of the end of the canal and the beginning of the acinar group. The functions 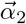 and 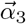 represent the left and right sides of the ductal group, respectively. Finally, the function 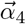 describes the lower part of the structure that encloses the ducts. In Fig. 9 it is shown schematically the geometry and definition of the functions 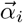, with *i* = 2, 3, 4. The geometry of the model allows us to calculate analytically the area in all regions and obtain that the domain area *A*_*T*_ is given by:

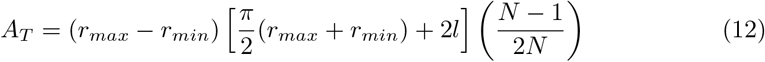

where N is the number of Voronoi cells in the domain and the last fraction represents the effective area distribution on the Voronoi diagrams.

**Fig 9.**
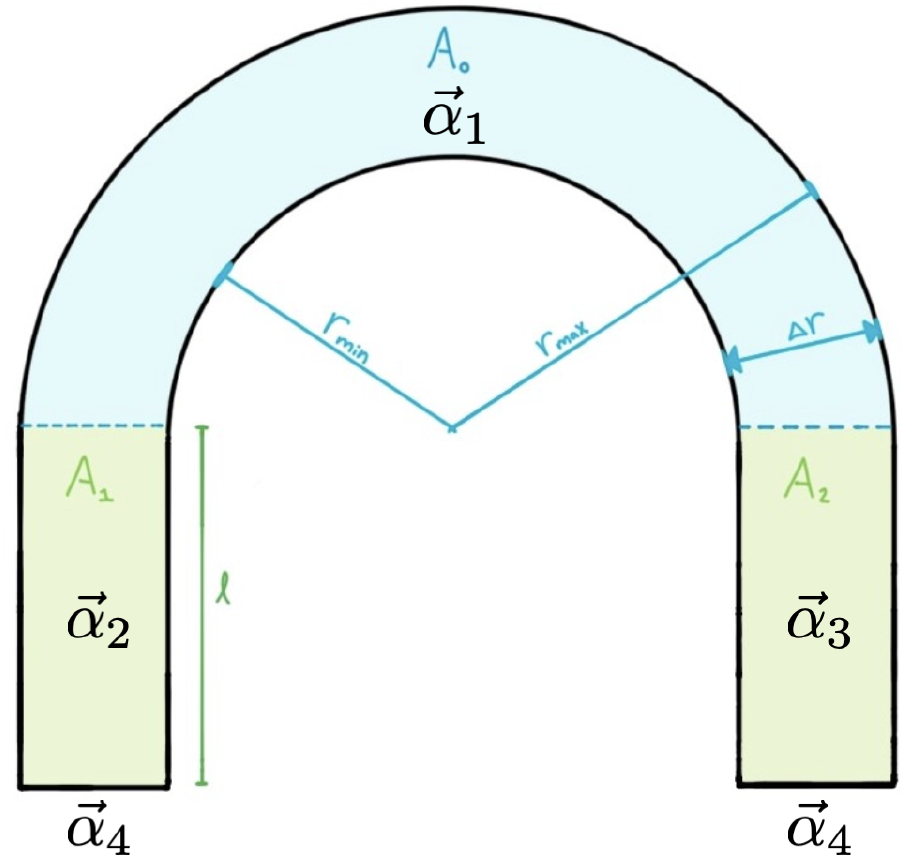
Domain configuration. *r*_*min*_ and *r*_*max*_ are the minimum and maximum radii, respectively. The functions 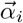 are defined in Eqs (11). *A*_0_ is the semicircle areas, *A*_1_ and *A*_2_ represent the left and right rectangular area.

### Anti-inflammatory mechanisms in the system

The acinar-ductal system undoubtedly contains mechanisms that allow it to mitigate inflammation and maintain an adequate balance in cellular tissue. Considering this observation, we believe that there are two key metrics in our model that can be used to evaluate inflammation in this system: (i) the effective area of the domain *A*_*T*_,and (ii) the basal cell size represented by its Voronoi area, *Â*_*cell*_. Thus, we define an inflammation indicator I by comparing the theoretical area for N cells at time t with the domain area, that is:

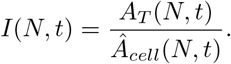

This indicator allows one to recognize situations in which the geometric domain introduces pressure on the cells (*I* = *A*_*T*_ /*Â*_*cell*_ > 1), which translates into an inflamed system because it does not have enough space to relax. On the other hand, when (*I* = *A*_*T*_ /*Â*_*cell*_ < 1), meaning that the cells have enough space to move and stabilize, that is, they are relaxed and even have extra space. In this sense, we propose two mechanisms to mitigate inflammation in the acinar-ductal system:

#### i) Elongation of the ductal canal depending on the numerical area and the area of the geometric domain

When there is growth, there is a decrease in both, the area as a function of cell density and perhaps the plasticity of the acinar-ductal system. Therefore, we proposed to lengthen the area of the ductal canal of the structure to reduce inflammation in the region and maintain an adequate balance in the tissue. This mechanism is activated when *I* = *A*_*T*_ /*Â*_*cell*_ > *A*_*up*_ is satisfied, where *A*_*up*_ is the threshold and is typically less than one. In our case, we have taken *A*_*up*_ = 0.9 which represents a 10% tolerance. Thus, the elongation of the ductal canal is calculated as follows:

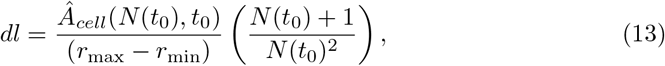

where dl is the length to be added to the ductal canal, *Â*_*cell*_(*N* (*t*_0_), *t*_0_) is the total area occupied by the cells at time *t*_0_, and *N*(*t*_0_) is the total number of cells.

#### ii) Fine adjustment of the length of the ductal canal for sudden changes in the number of cells

In the event that an accelerated increase in the number of cells occurs, an increase in the standard deviation of the area of the cells can be detected. To counteract this effect and mitigate inflammation, it is proposed to adjust the rate of change of ductal length. In this case, the activation mechanism occurs when *I* = *A*_*T*_ /Â_*cell*_ > *A*_*up*_ in order to ensure that cell density is conserved. In this case, the area change indicator Σ is defined as:

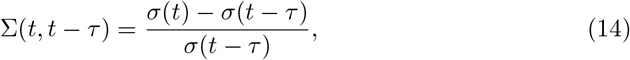

where Σ(t) is the standard deviation of the cell area at time *t*, and *τ* is a time interval that is related to the proliferation period. If the indicator Σ is outside the range given by the area increase thresholds, (Σ_min_, Σ_max_), an adjustment is made to the rate of change of ductal length. To maintaining a balance in the acinar-ductal system and mitigate inflammation in cellular tissue we propose the ansatz: Σ_min_ = −0.05 and Σ_max_ = 0.05 for the thresholds. To regulate cell density and preserve homeostasis in the tissue and/or considering sudden changes in the number of cells we adjusted the length of the duct. To do this, we consider two cases: a) *Inflammation*: Σ(*t, t* − *τ*) > Σ_max_, then the rate of change of the length *dl* of the acinar duct *dl* → *dl* + Σ(*t*) − Σ(*t* − *τ*); b) *Relaxation*: If Σ(*t, t* − *τ*) < Σ_min_, then the rate of change of the length dl of the acinar duct is decreased as *dl* → *dl* + 0.5 [*Σ(t)* − *Σ(t* − *τ*)].

### Gene functions

**Table 2.**
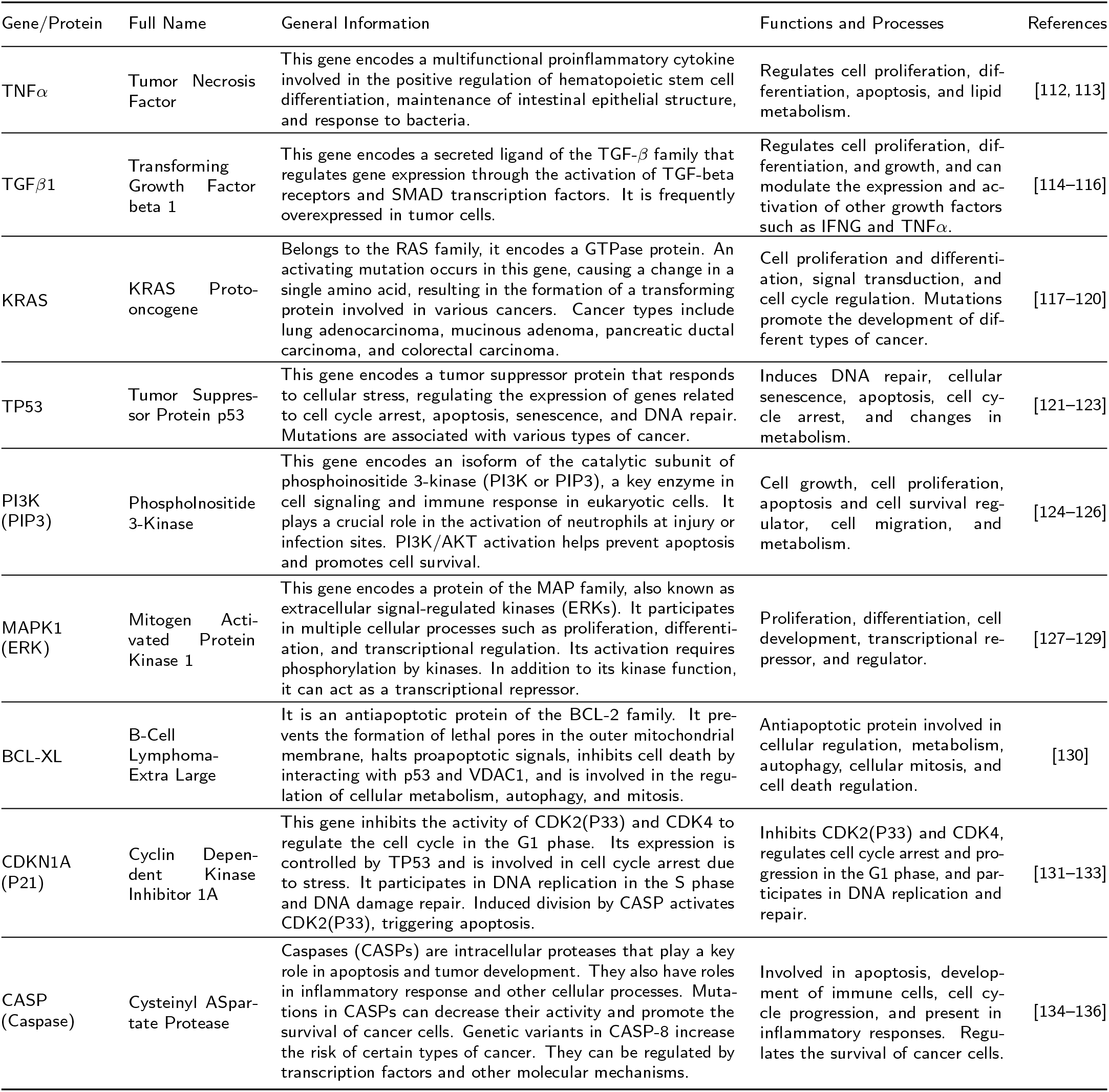
Overview of genes and cytokines in the reduced genetic regulatory network of Pancreatic Cancer.

## Acknowledgments

J.B.A. thanks the DGAPA of the UNAM for supporting the research and degree process through the scholarship awarded by the DGAPA-PAPIIT UNAM under grant number IN-109722. Likewise, he thanks to IIMAS and IMATE for their support in the development of this research. J.R.R.A. thanks the Department of Mathematics and Mechanics of the IIMAS and DGAPA-PAPIIT UNAM under grant number IA-100823. Also, J.R.R.A thanks to Ana Pérez Arteaga and Ramiro Chávez Tovar for their technical support. G.R.S. thanks the financial support of DGAPA-PAPIIT UNAM under grant number IN-109722.

